# Recovery Units under the Endangered Species Act could be used more widely

**DOI:** 10.1101/2020.03.15.991174

**Authors:** Michael Evans, Ya-Wei Li, Jacob Malcom

## Abstract

Recovering species is one of the main goals of the Endangered Species Act (ESA). In the face of limited budgets, diverse tools are needed to find efficient solutions. Recovery units may be one such tool - designated portions of a species’ range that must be recovered individually before an entire species can be considered recovered. Recovery units allow for spatial flexibility in recovery goals and may be used in regulatory decisions such as section 7 consultation. Despite their availability, there is very little information on how recovery units have been developed and used. We mined available public data to determine the number and types of species for which recovery units have been designated; evaluated species and geographic characteristics associated with recovery unit designation; and examined how recovery units have been used in the implementation of the ESA, such as during consultation. We found 49 listed species have designated recovery units through 2017, and that these species had similar characteristics. Namely, they had relatively large ranges and were well-studied. We found taxonomic biases in recovery unit designation as well, with fish species being disproportionately likely to have recovery units and plants disproportionately less. Improvements in recovery priority numbers among species with recovery units indicate that the theoretical benefits of this tool may have translated to improved status. These data indicate that recovery units could be applied to more wide-ranging species to improve recovery under the ESA.

## Introduction

Most species listed as threatened or endangered under the U.S. Endangered Species Act (ESA) are not yet recovered (Neel et al. 2012). The threats they face are increasingly diverse, and the agencies responsible for their recovery are challenged with limited budgets that do not match the growing number of listed species (Gerber 2016). Thus, conservationists need to develop and apply methods that can improve the effectiveness and efficiency of species recovery. Opportunities abound to do that through administrative reforms, without amending the ESA. The use of species “recovery units” is one such example of an existing tool within the ESA framework that may currently be underused.

Recovery units allow the agencies responsible for overseeing recovery under ESA – the U.S. Fish and Wildlife Service (FWS) and National Marine Fisheries Service (NMFS) – to split the range of a species into multiple units for recovery planning, rather than treating the species as a single entity. The Services’ Recovery Planning Guidance Handbook defines a recovery unit as “a special unit of the listed entity that is geographically or otherwise identifiable and is essential to the recovery of the entire listed entity” (National Marine Fisheries Service 2018). Thus, each recovery unit must be conserved to recover the species. The handbook also outlines how recovery actions and criteria may differ among units, allowing for more targeted and efficient recovery planning. Because the Services can delineate recovery units using a variety of factors – including genetic diversity, ecosystem diversity, and variation in threats – they can apply the tool to a wide range of taxa.

Section 7 of the ESA is one of the most important tools for protecting listed species (Malcom & Li 2015; Evans et al. 2019) and is particularly relevant to understanding the utility of recovery units. Section 7(a)(2) requires federal agencies to ensure that any actions they take, fund, or authorize do not jeopardize the existence of any listed species or destroy or adversely modify critical habitat. For species without recovery units, the Services evaluate whether a proposed federal action is likely to jeopardize the entire species. But for species with recovery units, the agencies may conduct the jeopardy analysis on the affected recovery unit. As the Services’ Section 7 Handbook explains, “when an action appreciably impairs or precludes the capability of a recovery unit from providing both the survival and recovery function assigned it, that action may represent jeopardy to the species.”

The Services rarely determine that actions jeopardize species or adversely modify critical habitat (Malcom & Li 2015; Evans et al. 2019), in part because most actions affect only a small fraction of a species’ range. Because FWS rarely tracks the cumulative effects of these actions (Office 2009), it may authorize levels of habitat disturbance that impede species recovery (Evans et al. 2016; Li et al. 2020). By evaluating the effects of proposed actions through the lens of recovery units, the Services may be more likely to determine these actions may jeopardize listed species. For example, if a species’ total range is 100 km^2^ and an action will remove 1 km^2^ of habitat, that 1% loss is unlikely to trigger a jeopardy conclusion. But if the same species has ten, 10 km^2^ recovery units and the same action affects one of those units, the loss is now 10% for the affected unit. That loss increases the risk of impairing the survival and recovery of the unit, which in turn increases the risk of jeopardizing the entire species. Thus, recovery units provide the Services with a tool to reduce the chances of a ‘death by 1,000 cuts’ scenario.

Here we present a series of analyses that advance our understanding of the Services’ use of recovery units through 2017 and evaluate the utility of recovery units for recovering species. Our first objective was to determine when and how the Services designate recovery units. Determining this fills an important gap, as no one has described how many or what types of species have recovery units. Our second objective was to assess how the Services use recovery units in recovery planning and section 7 consultations. Finally, we assessed whether species with recovery units show greater evidence of recovery than those without units. We use the results of these analyses to recommend how recovery units can be more effectively used to conserve listed species. None of these recommendations requires legislation, as the Services have all the necessary authority to implement the recommendations.

## Methods

All tabular data and code used to conduct analyses are available in an Open Science Framework repository, DOI 10.17605/OSF.IO/HNR46 (Evans 2019).

### Data for Recovery Unit Characteristics and Patterns

We used data from the Services to characterize and quantify patterns of recovery unit designation and use. Recovery units are designated in recovery plans, and recovery plans are written only for domestic U.S. listed species. We therefore considered only species with existing recovery plans (which is about 75% of ESA-listed species [Malcom & Li 2018]) in our analyses and refer to this set as “all species.” We obtained all available recovery plans from FWS’s ECOS website (https://ecos.fws.gov) and NMFS’s recovery site (http://www.nmfs.noaa.gov/pr/recovery/plans.htm). To identify the species with recovery units, we performed optical character recognition on all recovery plans, searched these documents for the term “recovery unit,” and then manually inspected matching documents to ensure true positives.

We considered several predictor variables that may explain why some species have recovery units and others do not. We collected species listing status (threatened or endangered), taxonomic membership, geographic region, range size, and recovery prioritization number (RPN) from the ECOS Recovery Plan Ad Hoc Report using the ‘ecosscraper’ package (https://github.com/jacob-ogre/ecosscraper) for R (R Core Team 2017). Ecosscraper is no longer maintained, and these data can now be obtained through the FWS Data Explorer API (https://ecos.fws.gov/ecp/report/ad-hoc-documentation?catalogId=species&reportId=species). Geographic region refers to the lead FWS Regional Office responsible for a listed species, or NMFS. To estimate range sizes, we obtained from ECOS the list of counties in which the Services report each species occurs. In addition to extracting the raw county data from ECOS, we identified ∼400 species that had ambiguous county occupancy data and manually refined the county lists using specimen and imagery databases (e.g., (GBIF [“GBIF: The Global Biodiversity Information Facility” 2020], eBird [Sullivan et al. 2009]). We joined these records with U.S. county data (Bureau 2019) to calculate the total area of occupied counties for each species. This overestimated the area occupied by each species because species typically occupy only select habitats in any county. For our analyses, we assumed the over-estimation did not bias towards species with or without recovery units.

The Services use RPNs to prioritize recovery efforts among listed species. The RPN scores range from 1 - 18, with 1 representing high priority, and are based hierarchically on the degree of threat a species faces (‘High’, ‘Moderate’, or ‘Low’), the species’ potential for recovery (‘High’, or ‘Low’), and its taxonomic uniqueness (‘Monotypic genus’, ‘Species’, ‘Subspecies’). Additionally, the Services may designate a species as potentially in conflict with economic activities using a ‘C’ suffix to RPNs (e.g., ‘2C’; [U.S. Fish and Wildlife Service 1983]. We extracted RPNs from the recovery plan table on ECOS and separated the priority number and conflict designation into two variables, Priority and Conflict.

Because the Recovery Handbook (National Marine Fisheries Service 2018) references the importance of genetic diversity and robustness for delineating recovery units, we considered the relative amount of genetic research for a species as a potential predictor of recovery unit designation. That is, a species that has been subject to a substantial body of genetics research may be more likely to have recovery units than a poorly studied species. We used Google Scholar to search for papers matching the term “[Species] population genetics,” and used the number of citations returned as a proxy for the extent of scientific knowledge of a species’ population genetics. We refer to this measure as “genetic citations.”

### Analyses of Recovery Unit Characteristics and Patterns

We hypothesized that species range sizes would differ among taxa. Similarly, we expected the rate of genetic citations to increase over time, due to growth of research in this field. We used one-way analysis of variance (ANOVA) to test for differences in species range size among taxonomic groups, and a linear model of the number of genetic citations among all species as a function of year between 1985 to 2015. If significant (α < 0.10) relationships were indicated for these measures, we converted species range sizes to standardized z-scores per taxonomic group, and genetic citation rates to standardized z-scores relative to the expected value for a given year. This allowed us to avoid confounding range size with taxonomic status, and level of genetic research with age of recovery plans when evaluating which species characteristics predict recovery unit designation.

We used a bootstrapping procedure to compare characteristics of species with recovery units to all species with recovery plans. We compared mean RPN, range size, genetic citations, and proportions of species designated as conflicting with development between a random sample of 49 species without recovery units to the 49 species with units. This procedure was repeated 100 times and we evaluated the proportion of random samples with means greater than species with recovery *P* < 0.05 to be significantly different. We then used logistic regression to test for differences in the proportions of species with recovery units as a function of these factors. We fit models predicting the log odds of recovery unit designation as a function of taxonomic group, FWS regional office, adjusted citation rate, adjusted area, and recovery prioritization. To assess the statistical significance of differences in species proportions we used Wald’s Chi-squared tests, and Tukey post-hoc tests for pairwise comparisons between specific taxa.

To characterize the species that have recovery units we applied a classification tree analysis (Hothorn et al. 2006) predicting recovery unit designation based on taxonomic group, adjusted citation rate, adjusted area, RPN, FWS regional office, and listing status. We used a minimum threshold of α < 0.90 for node creation and evaluated tree performance using receiver operating characteristic (ROC) curves. Because of the overrepresentation of plants among all U.S. listed species (∼57%), we generated trees using all species with recovery plans and using all species excluding plants, then selected the tree with the greatest predictive ability indicated by the area under the curve (AUC). We also used ROC curves to identify the appropriate class probability threshold for prediction, as the value maximizing the ratio of sensitivity to specificity. We generated classification trees and ROC curves using the *party* (Hothorn et al. 2006) and *pROC* (Robin et al. 2011) packages for R (R Core Team 2018).

We further investigated the relationship between species characteristics and recovery unit designation by comparing each species with recovery units to 1 - 3 taxonomically similar listed species without recovery units. We chose taxonomically similar listed species with recovery plans, prioritizing shared genera and no more distantly related than a shared family. For example, Preble’s meadow jumping mouse (*Zapus hudsonius preblei*) was compared to the New Mexico meadow jumping mouse (*Zapus hudsonius luteus*), and El Segundo blue butterfly (*Euphilotes battoides allyni*) was compared with Smith’s blue butterfly (*Euphilotes enoptes smithi*). We used conditional logistic regression (Connolly & Liang 1988) to estimate the log odds of recovery unit designation as a function of the same set of predictor variable used in classification tree analyses, eliminating taxonomic group. First, we fit univariate models for all predictors, and then a full model including all variables that were significant (*P* < 0.10) univariate predictors.

### Recovery Units in ESA Implementation

To understand how the Services use recovery units, we examined all recovery plans with recovery units for four criteria:

1. Does the recovery plan explicitly state that the recovery units are “essential to the recovery of the entire listed entity,” as explained in the Recovery Handbook?
2. On what basis did the Service designate the recovery units? We characterized the reasons according to themes in the Recovery Handbook, including variation in threats, and preservation of redundancy, representation, and resilience (‘3Rs’) related to geographic or genetic distinctiveness.
3. Does the recovery plan refer to the role of recovery units in section 7 consultations, as discussed in the Section 7 Handbook?
4. Does the recovery plan enumerate recovery criteria and/or actions for each recovery unit?

We determined that actions/criteria were enumerated per recovery unit only when differences among recovery unit were described in a recovery plan. Thus, if recovery units were referenced but actions were listed generically for all units (e.g., “High-quality habitat sufficient to ensure long-term survival and recovery is protected within each recovery unit”), we did not consider that recovery plan to have enumerated these factors per unit.

To assess the use of recovery units during section 7 consultation, we examined biological opinions (BiOps). For each species with recovery units, we randomly sampled up to ten BiOps from FWS’s TAILS database as replicated in Defenders of Wildlife’s Section 7 Explorer (Malcom and Li 2015; https://defenders-cci.org/app/section7_explorer/). We restricted the sampled population of consultations to those initiated after the Service published the recovery plan that designated the recovery units. We evaluated two aspects of each BiOp: whether species’ recovery units were mentioned and whether recovery units were used in the jeopardy analysis. We determined that recovery units were used in the analysis if one of two criteria were met:

1. The Service used recovery units as the basis for comparison when estimating the amount or effect of expected incidental take (harm to a listed species incidental to an otherwise lawful activity). Either number of species relative to the population within the recovery unit, or amount of habitat loss relative to the area of the unit.
2. The Service explicitly stated the role of recovery units in its jeopardy analysis.

We tested for differences in recovery unit mention and use in jeopardy analysis among taxa and FWS Regional Offices using Chi-square contingency tests. When comparing these rates among the Regional Offices, we collapsed records by BiOp, which can include multiple species. To determine whether the recency of recovery unit designation affected the probability that recovery units were used in BiOps, we modeled the odds of mention and usage as a function of days elapsed between a BiOp and designation (i.e., date of recovery plan) using logistic regression.

### Recovery Units and Recovery Progress

We examined five-year reviews to compare the recovery process of listed species with and without recovery units. At the most basic level, we determined the extent to which five-year reviews discuss recovery units. For every species with recovery units, we calculated the proportion of five-year reviews that mentioned the units among all reviews completed after the units were designated. We also determined how often the five-year reviews reported the status of species and recovery objectives by recovery unit. Five-year reviews also provide recommendations as to whether changes in listing status or recovery prioritization are warranted (i.e., recommended delisting, increase in priority number, or down listing from endangered to threatened). We recorded changes in species’ recovery priority numbers as a proximal indicator of status improvement, considering only changes reflecting either reduced threat level, or increased recovery potential. For example, a change from 2 to 3 represents a reclassification from species to subspecies, and not an improvement in status. Recommended down listings also indicated status improvement. We tested for differences in the proportion of species showing improvement at five-year reviews between species with recovery units and a random sample of 560 five-year reviews for all species with recovery plans. To evaluate whether these proportions were statistically different, we performed a bootstrapping procedure taking random samples of five-year reviews and comparing the frequency with which improvements were observed in each sample to the observed frequency among species with recovery units. We took 100 samples of size equal to the number of five-year reviews available for recovery unit species and used the proportion of samples with a higher frequency as our measure of significance.

## Results

### Recovery Unit Characteristics and Patterns

We identified 40 FWS recovery plans designating recovery units for 49 listed species (Table 1). We digitized the boundaries of all units that had maps, and the GIS files are available in a web map. The number of units per species ranged from 2 to 19. Units were as small as 7 acres and as large as 12,492,233 acres (Table 1). The rate of recovery unit designation has remained consistently low, between 0 and 3 species per year since 1995, except for eight species given recovery units in 2003 (Fig. 1).

**Table 1.**
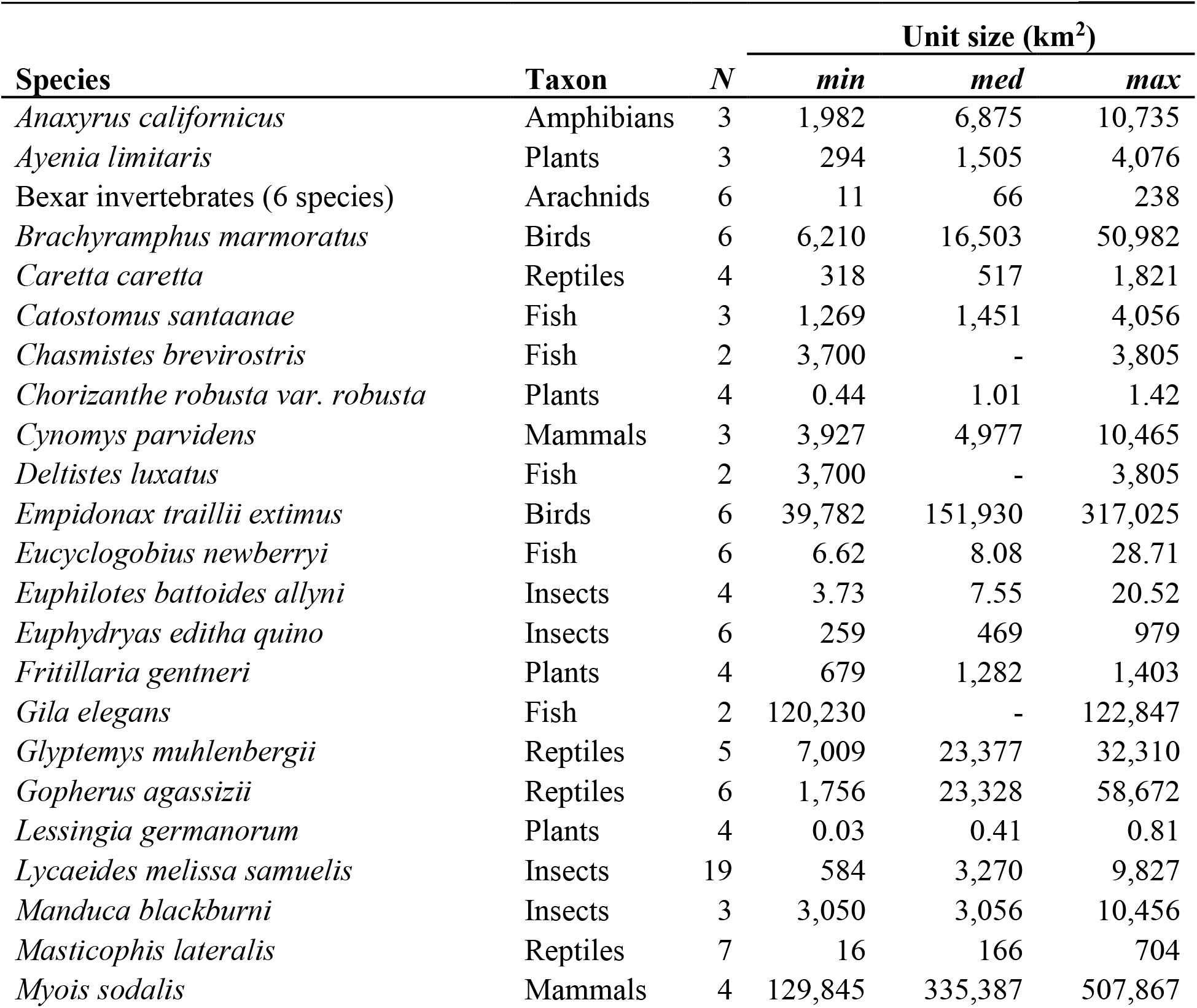
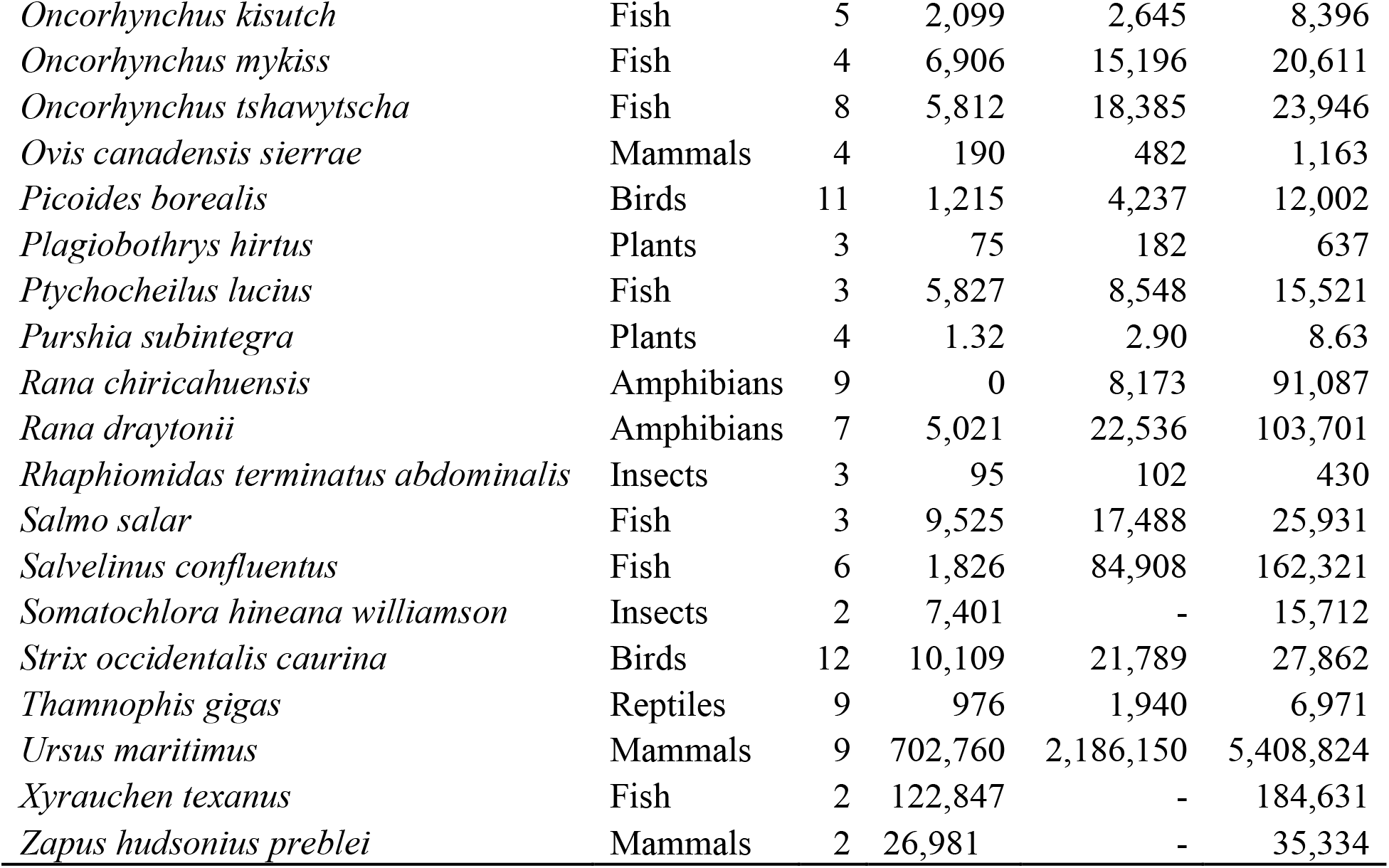
Species with recovery units designated as of as of December, 2017.

**Figure 1.**
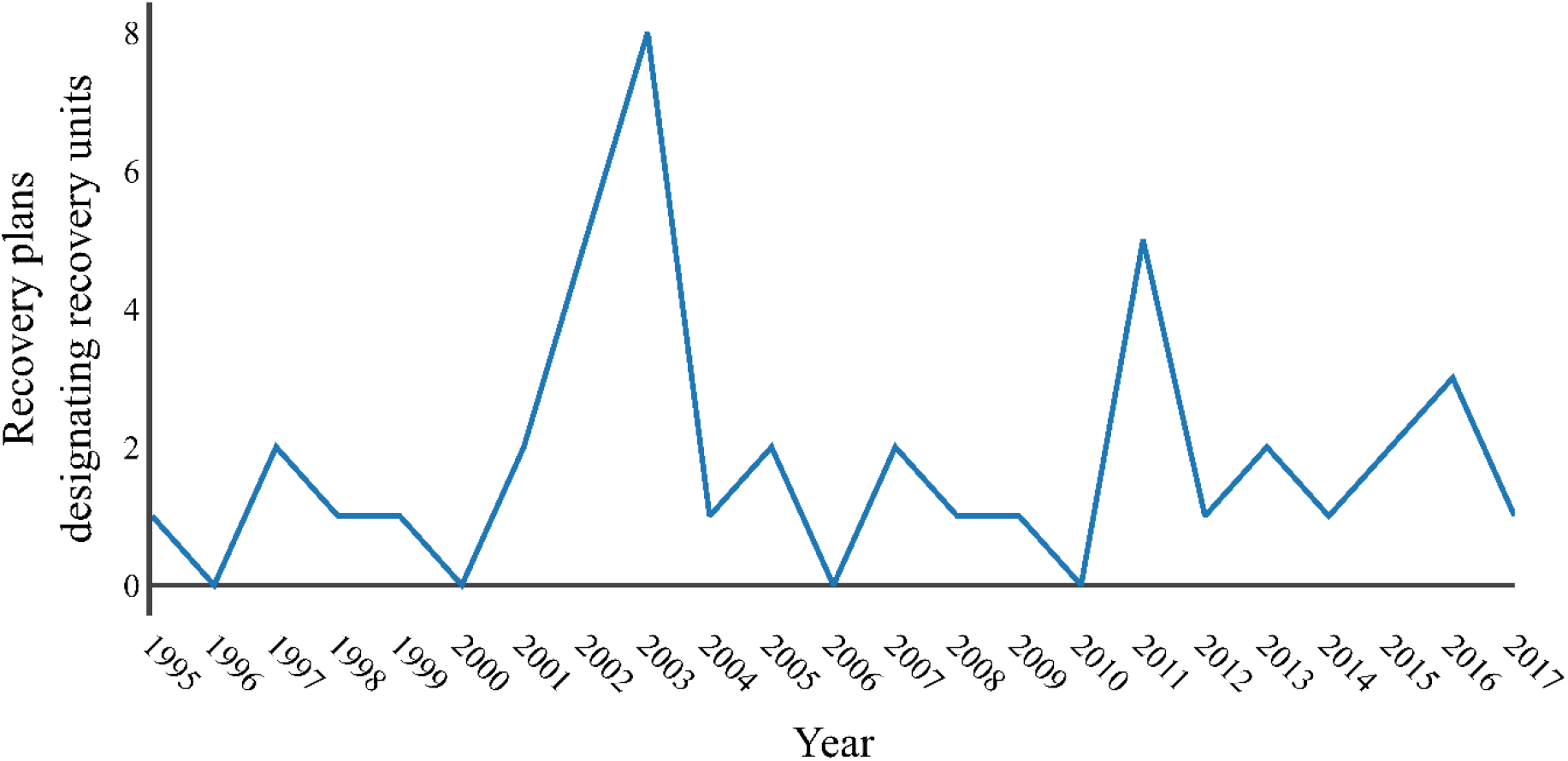
Recovery unit designation has been constant through time, except for 2003. Plot shows the number of recovery plans designating recovery units for threatened and endangered species per year from 1995 – 2017.

Species with recovery units tend to have larger range sizes than taxonomically similar species without recovery units. Species range sizes differed significantly among taxonomic groups (Table 1), as indicated by an ANOVA using the log of the area of occupied counties as the response variable (F_9,1352_ = 18.72, p < 0.001). Thus, we used standardized z-scores of areas per taxonomic group to account for differences in means among taxa when performing statistical tests using range size. Similarly, the number of genetic citations increased over time (β = 8.81 ± 1.25, *t* = 7.04, *P* < 0.001), and we transformed raw genetic citation numbers to z-scores per year. Bootstrapping procedures indicated taxonomically adjusted mean range size was significantly (*P* < 0.001) greater among species with recovery units (μ = 48,621,401 ac, s = 65,707,842 ac) than among all species with recovery plans (μ = 9,211,038 ac, s = 36,022,003 ac). Annually adjusted mean number of genetic citations were higher (*P* < 0.001) for recovery unit species (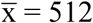, σ = 1127) than for all species with recovery plans (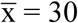, σ = 146.9), as shown in Fig. 2. Mean RPN did not differ between species with and without recovery units (*P* = 0.683). However, a significantly (*P* < 0.001) greater proportion of species with recovery units had an economic conflict designation (72%) than did all species with recovery plans (29%), and all listed species as of 2014 (27%).

**Figure 2.**
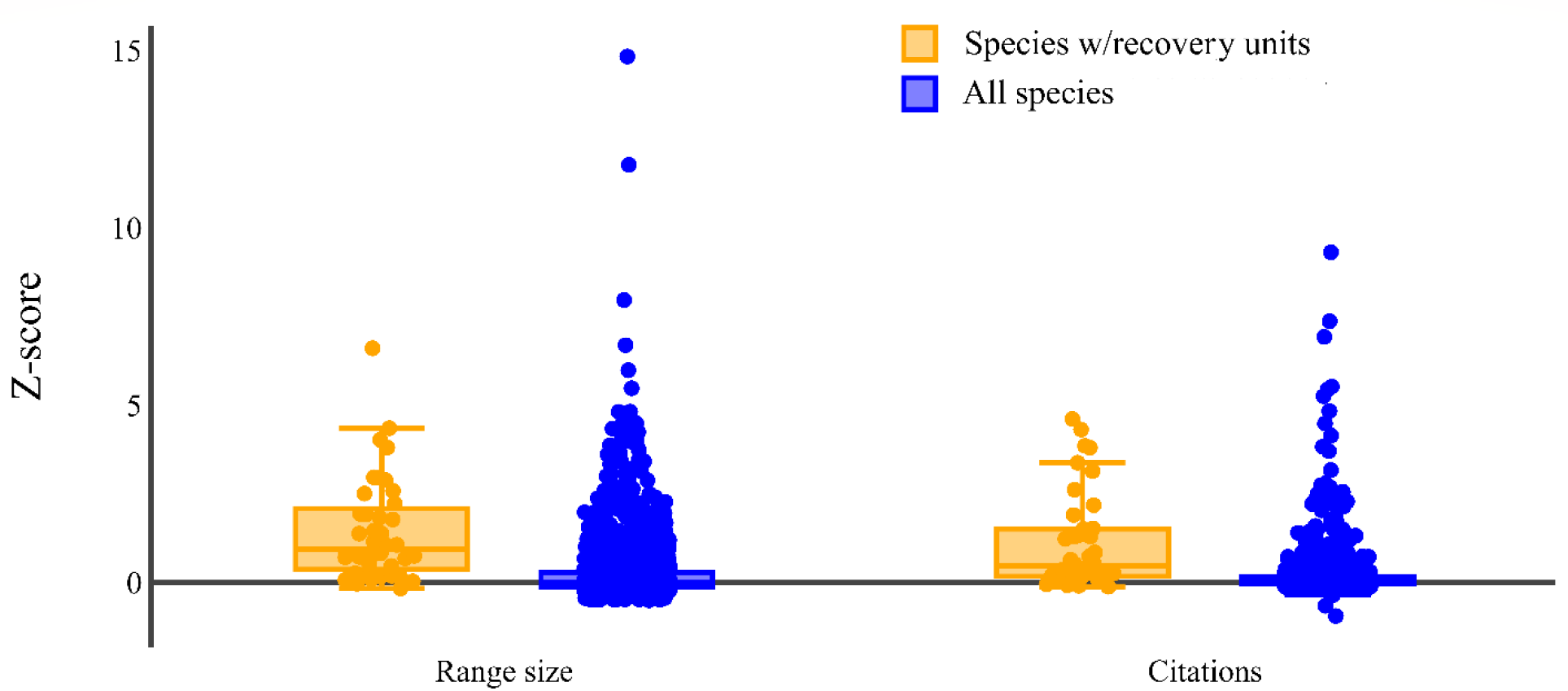
Species with recovery units have larger range sizes and a greater amount of published genetics research, on average, relative to all listed species. Standardized range sizes and number of Google Scholar citations including the term ‘population genetics’ were higher for species with recovery units (orange) than all species with recovery plans (blue).

Designation of recovery units differed significantly among taxa (X^2^ = 101.43, df = 9, p < 0.001). Amphibians, birds, fish, insects, mammals, and reptiles are more frequently given recovery units relative to their frequency among listed species with recovery plans (Fig. 3). The odds of designation for plants were significantly lower than those for amphibians (*Odds Ratio* = 0.069, *P* = 0.006), fish (*OR* = 0.082, p < 0.001), insects (*OR* = 0.085, *P* < 0.001), mammals (*OR* = 0.101, *P* = 0.005), and reptiles (*OR* = 0.043, *P* < 0.001). No other odds ratios were significant. There were no differences in frequencies of recovery unit designation among species between FWS regions (X^2^ = 48, df = 42, *P* = 0.243).

**Figure 3.**
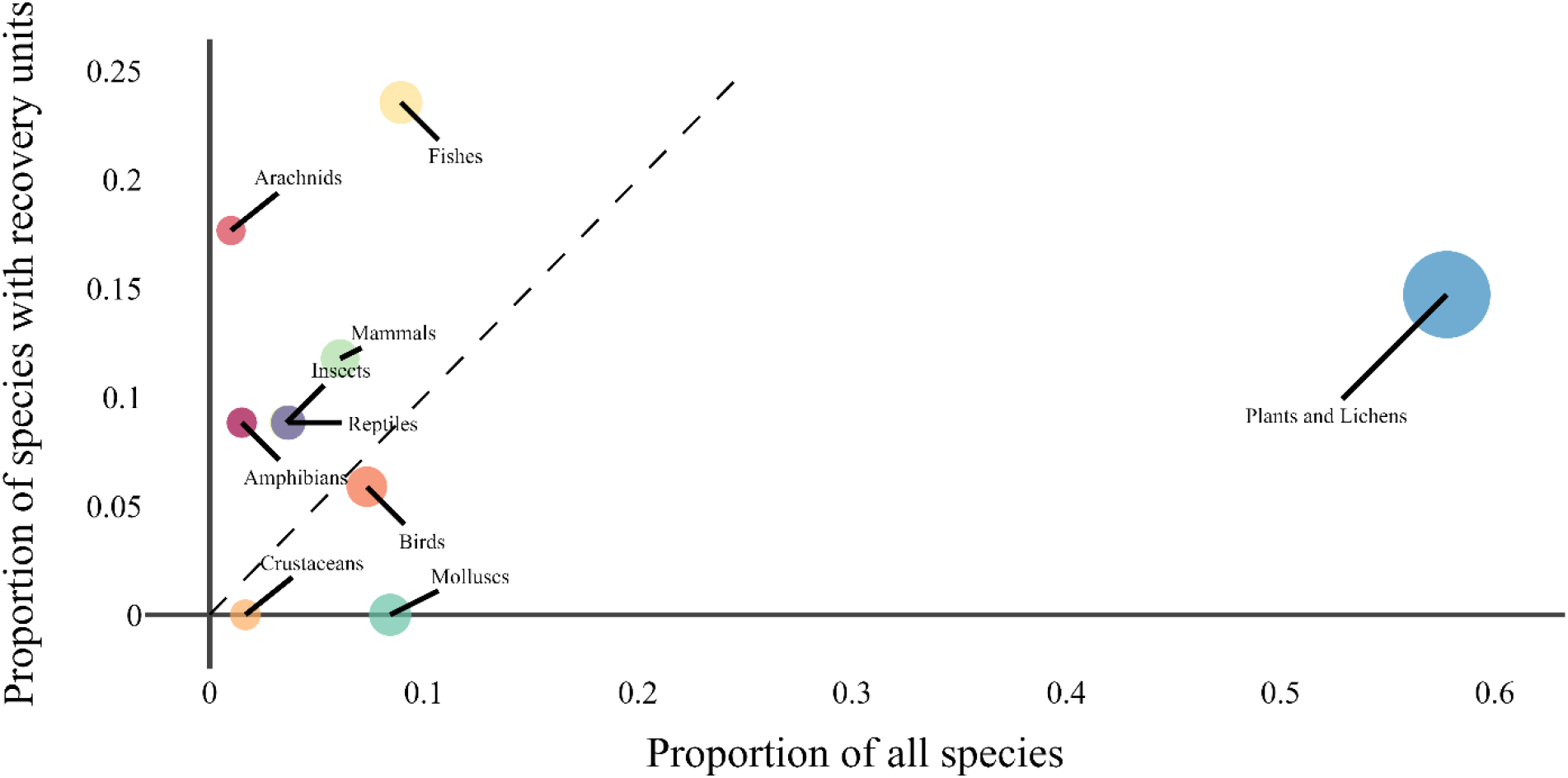
Designation of recovery units is biased among taxonomic groups. The dotted line indicates an expected 1:1 relationship if the probability of recovery unit designation did not depend on a species’ taxonomic grouping. Fishes received disproportionately more designations than expected, whereas plant species received disproportionately fewer.

Classification trees built using all species with recovery plans exhibited better predictive performance of recovery unit designation (AUC = 0.91) than trees build excluding plant species (AUC = 0.84). The best performing tree included FWS office, taxonomic group, citation rate, and range size (Fig. 4a). When FWS office was excluded, the resulting tree indicated that species with range sizes above the 71st percentile of their taxonomic group, and annually adjusted genetic citation rates above the 72nd percentile had a 0.70 probability of having recovery units designated. Taxonomic group and FWS office were important predictors for species falling below the range size and citation rate thresholds (Fig. 4b). We identified a classification probability threshold of 0.467 as providing the maximum ratio of sensitivity and specificity.

**Figure 4.**
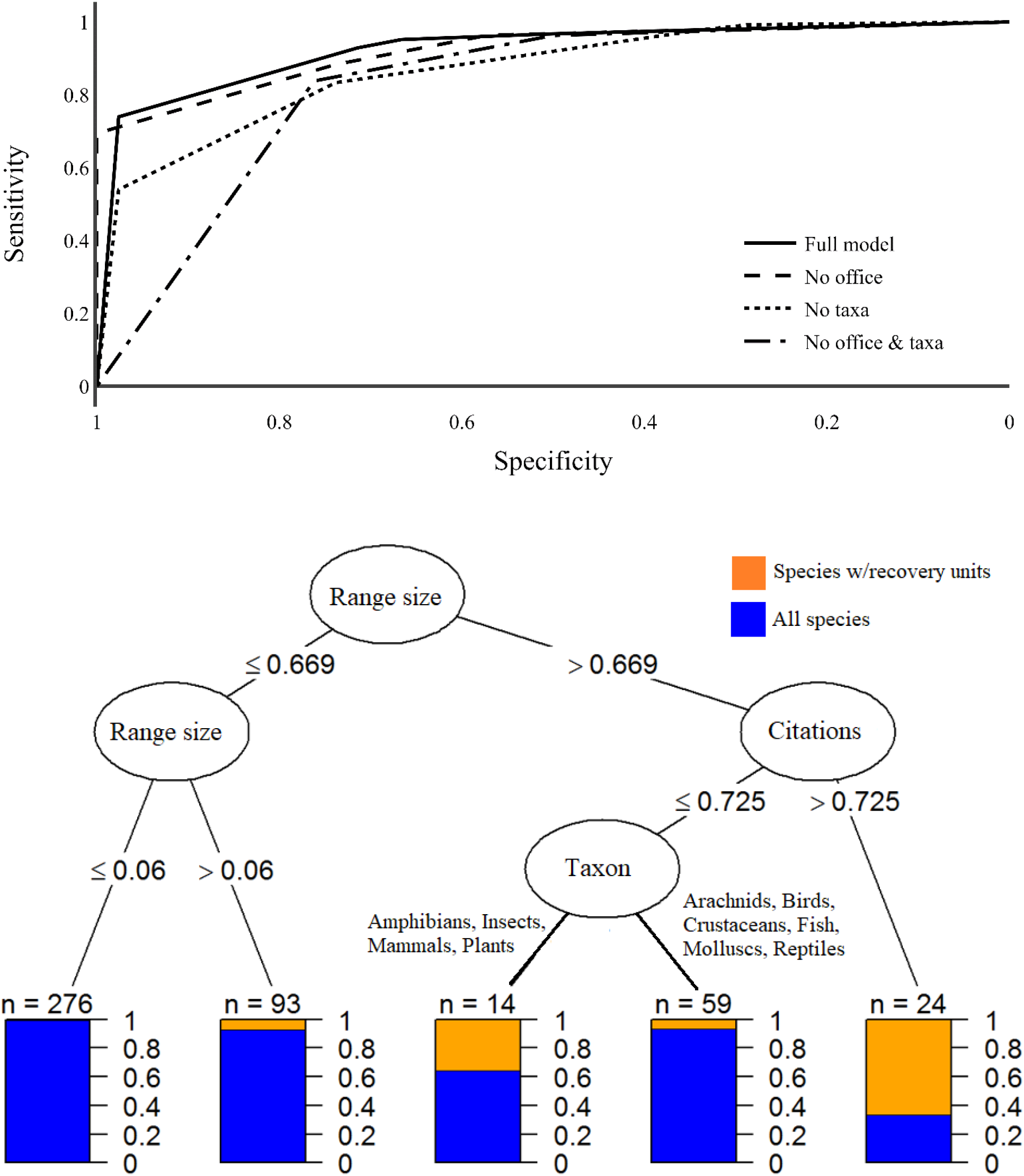
A classification model considering all variables was most effective at predicting recovery unit designation. Removing FWS office from the classification model lowered predictive ability negligibly, as indicated by the area under the curve (upper). Results from a model excluding FWS office shows effects of species range size and genetic citation rate predicting species with recovery units (lower). Values shown along edges are standardized z-scores within taxonomic groups for range size, and per year for genetic citations.

Results from conditional logistic regression corroborated results from classification tree analysis. The only significant univariate predictors of recovery unit designation were genetic citations (*B* = 0.68 +-0.29, p = 0.019) and range size (*B* = 0.53 +-0.21, p = 0.011). Greater number of genetic citations and larger range size increased the probability of recovery unit designation. A full model including these predictors did not indicate any significant relationships between species characteristics and probability of recovery unit designation.

### Recovery Units in ESA Implementation

Of the 40 recovery plans designating recovery units, 24 stated that recovery units are ‘essential for species recovery’ (Table 2). Except for five species (*Lessingia germanorum, Masticophis lateralis euryxanthus, Ptychocheilus lucius, Somatochlora hineana williamson, Oncorhynchus kisutch*), all plans provided some explanation for the designation of recovery units in terms of their role and importance in facilitating persistence and recovery of the entire species. Explanations fell into two major categories: addressing variation in the threats species face (and needed recovery actions) between units; and addressing the ‘3Rs’ of conservation (Redundancy, Representation, and Resilience [Shaffer & Stein 2000; Wolf et al. 2015]). Of the 35 plans providing explanations, 29 plans referenced the importance of preserving either geographic and/or genetic variability and representation. All but 9 recovery plans specified different recovery actions and/or criteria per unit. Additionally, 10 recovery plans provided guidance on the role of recovery units during section 7 consultation, explicitly referring to the use of recovery units in jeopardy analysis (Table 2).

**Table 2.** Summary of details provided in recovery plans designating recovery units related to ESA implementation. Data include whether a plan described the role of units in jeopardy analyses (Jeopardy), whether the role of units as essential to species recovery is acknowledges (Essential), how the designation of units is justified (Justification), and whether unique recovery actions or criteria were specified for each recover unit (Actions). Page numbers are provided where each element occurred in a recovery plan.

| Species | Date | Jeopardy | Essential | Justification | Actions |
| --- | --- | --- | --- | --- | --- |
| *Anaxyrus californicus* | 7/24/1999 | NA | p77 | 3Rs (p75), Threats (p75) | p75 |
| *Ayenia limitaris* | 8/5/2016 | NA | p39 | 3Rs (p39) | NA |
| *Brachyramphus marmoratus* | 9/24/1997 | NA | NA | 3Rs (p115) | p125 |
| *Caretta caretta* | 12/31/2008 | NA | pII-2 | 3Rs (pII-2) | pix |
| *Catostomus santaanae* | 2/16/2017 | NA | pII-3 | 3Rs (pII-3) | pIII-5 |
| *Chorizanthe robusta robusta* | 12/20/2004 | NA | NA | 3Rs (p36) | p41 |
| *Cicurina barnia*, *C. madla*,<br>*C. venii*, *C. vespera*,<br>*Neoleptoneta microps*,<br>*Texella cokendolpheri* | 9/12/2011 | NA | p18 | 3Rs (p18) | NA |
| *Clemmys muhlenbergii* | 5/15/2001 | NA | NA | Threats (p31) | p42 |
| *Cynomys parvidens* | 3/1/2012 | p3.2-1 | p1.3-7 | 3Rs (p3.2-1) | NA |
| *Deltistes luxatus*,<br>*Chasmistes brevirostris* | 1/12/2013 | NA | NA | 3Rs (p39) | p45 |
| *Empidonax traillii extimus* | 3/8/2002 | NA | NA | Threats (p3) | p84 |
| *Eucyclogobius newberryi* | 12/7/2005 | NA | NA | 3Rs (p30) | p41 |
| *Euphilotes battoides allyni* | 9/28/1998 | NA | NA | 3Rs (p23) | NA |
| *Euphydryas editha quino* | 8/11/2003 | NA | p76 | 3Rs (vi), Threats (vi) | pvi |
| *Fritillaria gentneri* | 8/28/2003 | p39 | p34 | 3Rs (p16) | NA |
| *Gila elegans* | 8/1/2002 | NA | p6 | Threats (p6) | p42 |
| *Gopherus agassizii* | 5/6/2011 | NA | p41 | 3Rs (p41) | p72 |
| *Lessingia germanorum* | 8/8/2003 | NA | NA | NA | p128 |
| *Lycæides melissa samuelis* | 8/25/2003 | NA | NA | 3Rs (52) | p56 |
| *Manduca blackburni* | 9/28/2005 | p39 | p39 | 3Rs (p40) | pvi |
| *Masticophis lateralis euryxanthus* | 9/1/2002 | NA | NA | NA | pII-80 |
| *Myotis sodalis* | 4/16/2007 | NA | p116 | 3Rs (p8) | p124 |
| *Oncorhynchus gilae* | 9/10/2003 | NA | NA | 3Rs (p38) | p41 |
| *Oncorhynchus kisutch* | 9/1/2012 | NA | NA | NA | NA |
| *Oncorhynchus mykiss*,<br>*Oncorhynchus tshawytscha* | 7/1/2014 | NA | p68 | 3Rs (iv) | p96 |
| *Ovis canadensis sierrae* | 9/24/2007 | NA | p45 | 3Rs (p43) | pvi |
| *Picoides borealis* | 1/27/2003 | p147 | pxii | 3Rs (pxii) | pxv |
| *Plagiobothrys hirtus* | 9/25/2003 | p19 | p19 | 3Rs (p5) | p24 |
| *Ptychocheilus lucius* | 8/1/2002 | NA | NA | NA | p34 |
| *Purshia subintegra* | 6/16/1995 | NA | p1 | 3Rs (p51), Threats (p1) | p55 |
| *Rana chiricahuensis* | 3/14/2007 | p55 | p49 | 3Rs (p54), Threats (p54) | p57 |
| *Rana draytonii* | 5/28/2002 | p48 | p48 | Threats (p45) | p62, p79 |
| *Rhaphiomidas terminatus abdominalis* | 9/14/1997 | NA | NA | Threats (p10) | p17 |
| *Salmo salar* | 1/31/2019 | p4 | NA | 3Rs (p2) | NA |
| *Salvelinus confluentus* | 9/28/2015 | p176 | p33 | 3Rs (p33) | Sep. docs* |
| *Somatochlora hineana williamson* | 9/27/2001 | NA | NA | NA | p39 |
| *Strix occidentalis caurina* | 6/28/2011 | pIII-I | pIII-I | 3Rs (pIII-I) | NA |
| *Ursus maritimus* | 1/11/2017 | N | p25 | 3Rs (p25), Threats (p25) | NA |
| *Xyrauchen texanus* | 8/1/2002 | N | p6 | Threats (p6) | p36 |
| *Zapus hudsonius preblei* | 9/18/2018 | N | p40 | 3Rs (p40) | p55 |

\*Actions for bull trout recovery units provided in separate documents per unit

Generalized linear models indicated that species with recovery units also had a significantly higher mean number of section 7 consultations and rate of formal consultation (μ = 0.36, s = 0.26) than all listed species (μ = 0.24, s = 0.30), after accounting for differences between taxonomic groups, FWS offices, recovery priority number, listing status, and range size (F_800,799_ = 7.54, p < 0.01). Among these covariates, only taxonomic group, FWS office, and range size were also significant predictors of consultation rates. We read 216 BiOps that could have considered RUs in jeopardy determinations. Of these, 62% mentioned the existence of recovery units for the relevant species. Of those BiOps, 67% used recovery units in the jeopardy analysis. Overall, 42% of BiOps that could have used recovery units in the jeopardy analysis did so explicitly. These rates varied by taxa (X^2^ = 44.17, df = 7, p < 0.001) and FWS office (X^2^ = 94.47, df = 31, p < 0.001).

A post-hoc test for correlation indicated that species with high probability of recovery unit designation were also more likely to have their recovery units mentioned and used during section 7 consultation (Fig. 5). The odds of RUs being mentioned in BiOps decreased significantly (*B* = −3.24*10^−4^ ± 7.92*10^−5^, p < 0.001) with time. On average, the odds of mention drop to 1 (i.e., 50% chance) after 12 years. The odds that RUs were used in jeopardy analysis also decreased significantly (*B* = −2.153*10^−4^ ± 7.48*10^−5^, p = 0.004) over time, dropping to 1 after 5 years on average (Fig. 6).

**Figure 5.**
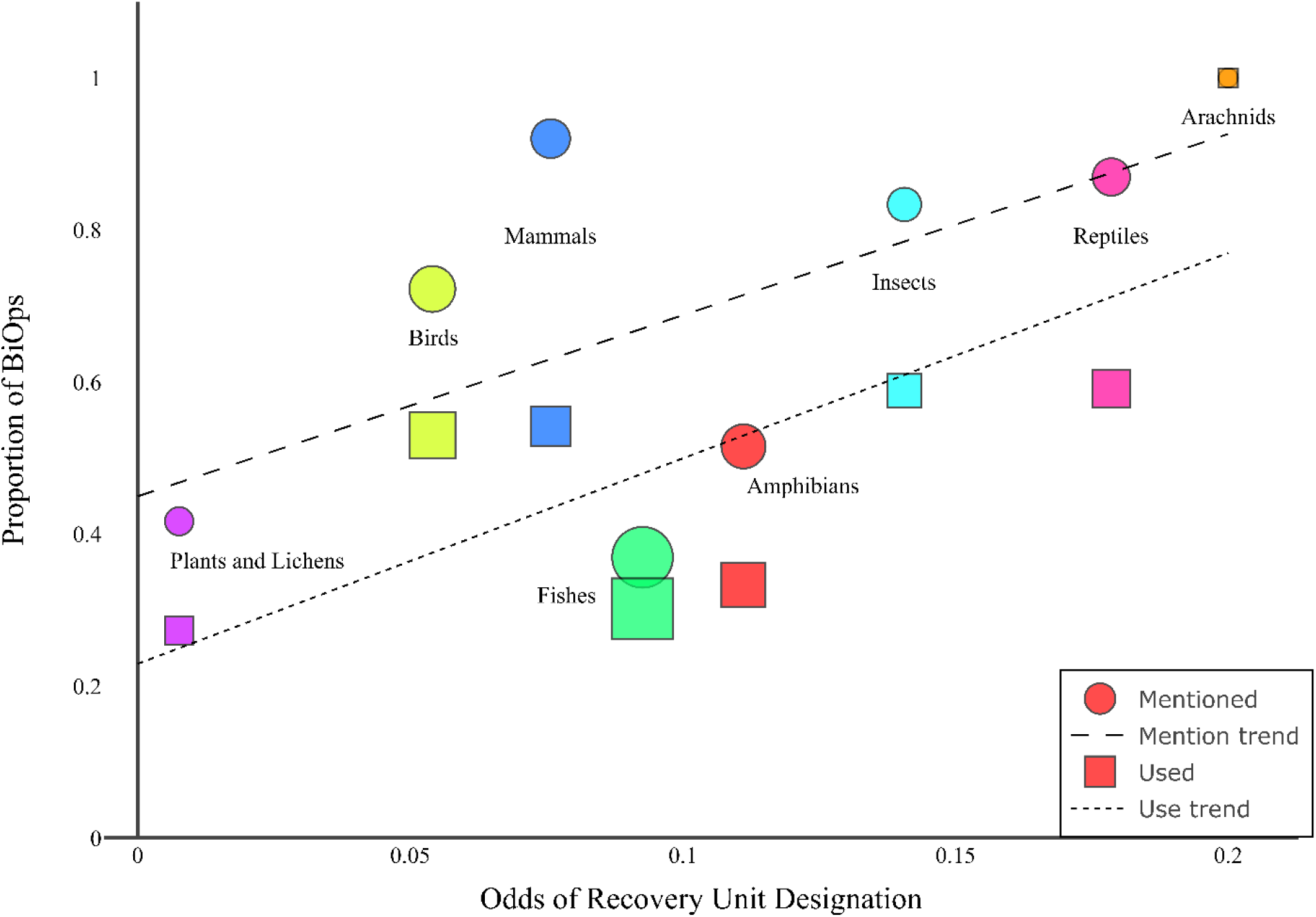
Taxonomic groups that were more likely to have recovery units designated were also more likely to have those units mentioned and used during consultation. Plot displays the proportion of biological opinions (BiOps) in which recovery units were mentioned or used as a function of the odds that recovery units were designated per taxonomic group. Marker size is proportional to the number of BiOps evaluated for each taxonomic group. The linear relationship between taxon odds of unit designation and recovery unit mention (dashed line), and use (upper line) in BiOps.

**Figure 6.**
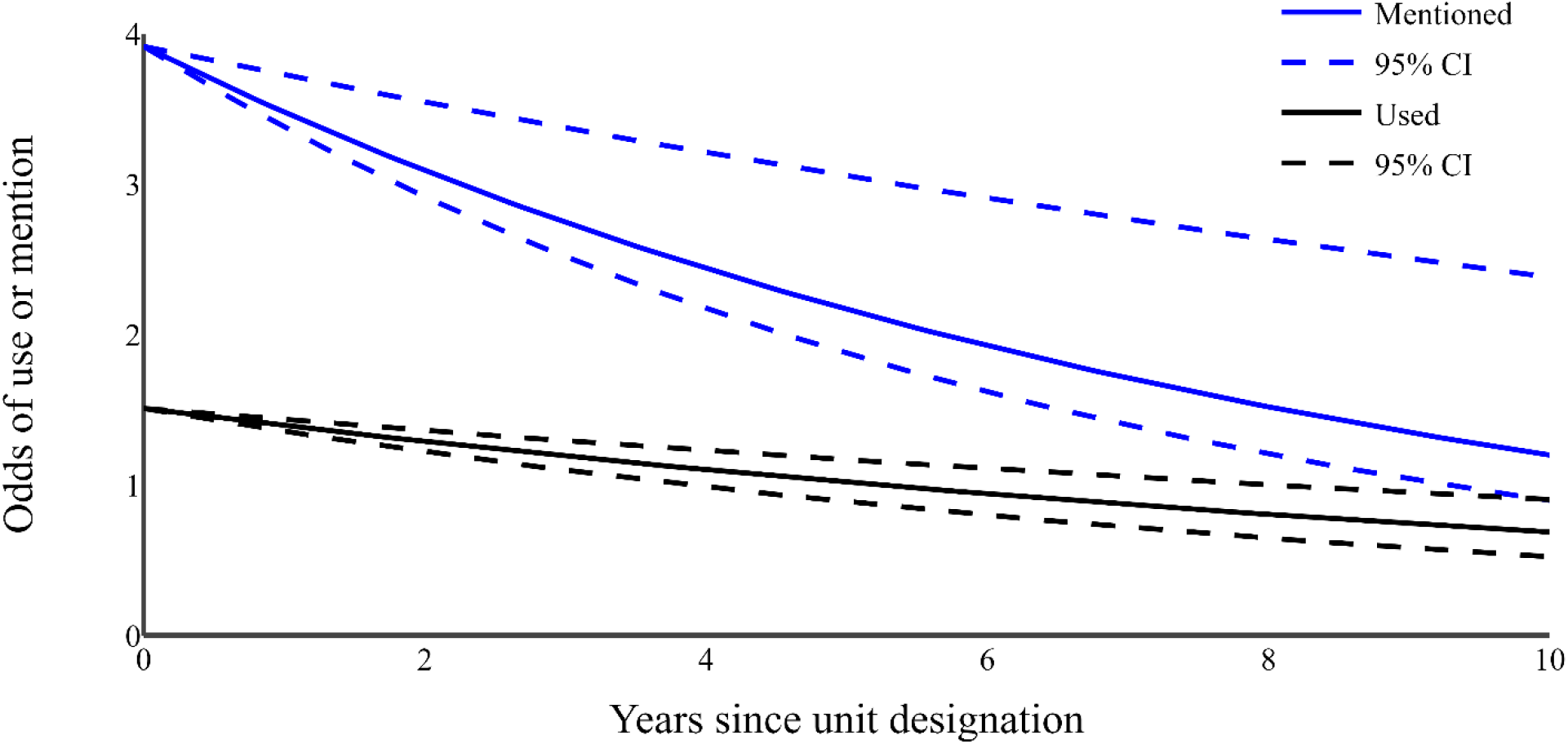
Recovery units were mentioned and used less often in ESA consultation over time. Curves show the odds that recovery units were mentioned and used during Sec. 7 consultation over time, as estimated by logistic regression. Confidence intervals around estimated slopes are indicated by dotted lines.

### Recovery Units and Recovery Progress

Of the 49 species with recovery units, 24 had five-year reviews conducted after the recovery plan designating the units was written. Of these 24 reviews, RUs were explicitly mentioned for all but one species (*Manduca blackburnii*). For these species, population statuses and recovery criteria were evaluated and reported by recovery unit. The frequency of species improvement, indicated by changes in recovery priority number, was significantly higher (p < 0.01) among species with recovery units (0.174), than the rate observed among all five-year reviews (0.086), as determined by bootstrapping.

## Discussion

Recovery units are an existing tool under the ESA that provide flexibility and potentially stronger conservation measures for listed species. Units can be used to refine recovery criteria within a species’ range, and in theory better align jeopardy analyses during section 7 consultations with the recovery goals for a species. Here we analyzed recovery plans, section 7 consultations, and five-year reviews to characterize how recovery units are designated and used under the ESA. Recovery units were designated for only 2.9% of U.S. listed species (as of January 2018), and our analyses indicated common characteristics among species receiving recovery unit designation. Species with a greater number of genetic studies and larger range size, were more likely to receive recovery units (Fig. 4). These criteria may not have been used explicitly by Service biologists during recovery plan development, but rather characterize well-studied, wide-ranging species that are inherently more likely to match the description (i.e., ‘geographic subsets’) and purpose (i.e., ‘preserve genetic robustness’) of recovery units in the Recovery Handbook. These patterns indicate the influence of guidelines presented in the recovery planning document. Taxonomic biases, as well as spatial and temporal variation in the designation and consideration of recovery units indicate opportunities to improve standardization in how this tool is used. Conceptually, and some empirical evidence suggest benefits of using recovery units, so rectifying these inconsistencies could improve species recovery under the ESA.

We found some evidence that the designation of recovery units corresponds to improved recovery relative to species without recovery units. Greater frequency of improvement in recovery priority numbers suggests species with recovery units exhibit either greater demographic improvement or reduction in threats. Recovery priority numbers are an imperfect means by which to assess species conservation status (Malcom et al. 2016), and these inferences are limited by the coarse nature of RPNs for assessing species status. However, they were the only dataset available providing an extensive, consistent means by which to quantify listed species recovery. Quantitative population data and monitoring reports, such as those provided in the 2012 Utah prairie dog five-year review, or a consistent key for scoring threat and demography status, would allow for more robust assessment of the effect on both recovery units and listed species recovery (Malcom et al. 2016).

If there is some benefit to recovery unit designation, then objective criteria for additional designation could help extend this benefit to other applicable species. The 491 species without a final recovery plan as of January 2018 provide perhaps the best opportunity to designate recovery units for appropriate species. Thresholds in important characteristics identified by classification tree analyses can be used to identify species that would be consistent with those for which the Services have historically provided recovery units. Our results showed that species in the upper 30th percentiles of range size relative to taxonomic means, and upper 27th percentile of annually adjusted genetic citation rate would be consistent with current recovery unit designation patterns. These results do not allow for an inference of causation, and it is unclear whether these criteria represent a best practice. Thus, many additional species not fitting this profile might also benefit from recovery units.

Patterns in recovery unit designation suggest variation in practices within the Services affected recovery unit creation and implementation. We found significant differences among FWS offices in the rate of recovery unit designation and their inclusion in BiOps. This variation was corroborated by our interviews with FWS staff who expressed differing views on recovery units. For instance, some staff hesitated to designate units because doing so may impede the delisting of species due to recovery if all but one unit has met recovery criteria. Others thought recovery units should be applied liberally. Many FWS staff noted that the completion of recovery planning training and the expertise of the species’ recovery team made it more likely that units were used. Personnel-specific expertise could explain the decrease in per species recovery unit acknowledgement in BiOps over time (Fig. 6), as an initial emphasis on the use of recovery units fades as new staff become involved with a species. Together, these patterns in space and time suggest a need for standardized practices and established institutional knowledge regarding the use of recovery units.

Overlooking recovery units during consultations could undermine species recovery, as consultations are one of the primary ways the ESA protects listed species. The ability to consider how an action affects a species at a refined geographic scale during consultation is one of the primary benefits that recovery units provide. However, some FWS staff expressed concern that recovery units would force biologists to call jeopardy, reflecting a lack of clarity regarding their implications. Other FWS staff doubted that recovery units would ever actually be used as the basis for a jeopardy finding. This observation makes sense for some extremely wide-ranging species (e.g., Northern spotted owl) because each recovery unit still covers a large enough area such that the great majority of federal actions are not extensive enough to seriously affect the entire unit. However, such scenarios do not explain the observed low proportion of BiOps that evaluated the effects of a proposed action at the recovery unit level (Fig. 5). For recovery units to be used to improve protection and recovery of listed species, FWS will at a minimum need to more clearly and consistently train staff on their designation and use.

One way to increase the application of recovery units would be for the Services to more frequently emphasize in recovery plans the use of recovery units as the unit of jeopardy analysis during consultations. Although jeopardy has rarely been called during consultations, the Services are able to use this process to negotiate conservation measures with regulated entities to avoid jeopardy (Malcom & Li 2015, Evans et al. 2019). Even absent a jeopardy determination, the smaller denominator provided by RUs may enable more conservation benefits. Considering that RUs were inconsistently used during section 7 consultation (Fig. 6), the impact of recovery units on recovery could potentially be even greater than observed.

Our examination of recovery plans indicated that the Services generally provide thorough and robust explanations for the designation and importance of recovery units. However, we found that only 10 out of 40 plans made the connection to jeopardy analysis explicit (Table 2). Recovery plans most often cited the importance of maintaining multiple sub-segments of a species’ population to preserve diversity and provide resilience, and FWS staff that supported the use of recovery units expressed similar reasons to designate recovery units. These explanations for using and delineating recovery units closely matched the reasons in the Recovery Handbook (e.g. ‘genetic robustness’, ‘demographic robustness’, ‘important life-history stages’). Thus, it seems the explanation for designating recovery units is important if they are to be used by the Services to uphold stronger protection for species.

While the guidance provided in the handbook leaves room for interpretation with the phrase ‘or some other feature necessary for long-term sustainability of the entire listed entity,’ it seems that the Services primarily use the specific examples identified. This presents a potential opportunity to expand the use of recovery units to offer more robust protection. For instance, population fragmentation (Crooks et al 2017) and climate change (Thomas et al 2004; Diaz et al 2019) are two of the most often cited threats to species persistence, aside from direct habitat loss. As connectivity and the capacity to adapt to climate change are clearly scientifically supported as necessary for long-term persistence, the Services might use recovery units to afford extra protection in areas of a species’ range providing connectivity and future capacity for range shifts.

Finally, we provide an important step towards achieving consistent implementation of recovery units by publishing spatially referenced GIS data for recovery units (Evans & Moskwik 2017). To our knowledge, unit maps exist only as static images in recovery plans. Accessible, geocoded maps can make it easier for the Services to consider recovery units during a jeopardy analysis. Currently, an inability to locate action areas within recovery units may contribute to the disparity between the rate at which recovery units are mentioned in BiOps and the rate at which they are used in jeopardy analysis, as spatial data is more essential to the latter (Fig. 5). GIS data for recovery units also provides a critical basis for further analyses investigating the potential role of recovery units in ESA implementation and species recovery. For example, variation in a species’ vulnerability or stability among recovery units can be informed by analyzing the distribution of public versus private land ownership, and critical habitat between units. Additionally, the distribution of spatially referenced Section 7 consultation locations among units could inform recommendations for adjusted levels of authorized take and disturbance on a per unit basis. These kinds of analyses are important to evaluate the utility of recovery units, and how they can continue to be used to improve endangered species conservation.

## Acknowledgements

We thank Services personnel for their time talking us through their views on and approaches to using recovery units in ESA implementation.

